# Study of the genetic and phenotypic variation among wild and cultivated clary sages provides interesting avenues for breeding programs of a perfume, medicinal and aromatic plant

**DOI:** 10.1101/2020.11.22.393264

**Authors:** Camille Chalvin, Stéphanie Drevensek, Christel Chollet, Françoise Gilard, Edita M. Šolić, Michel Dron, Abdelhafid Bendahmane, Adnane Boualem, Amandine Cornille

## Abstract

A road-map of the genetic and phenotypic diversities in both crops and their wild-related species can help identifying valuable genetic resources for further crop breeding. The clary sage (*Salvia sclarea L.*), a perfume, medicinal and aromatic plant, is used for sclareol production and ornamental purposes. Despite its wide use in the field of cosmetics, the phenotypic and genetic diversity of wild and cultivated clary sage remains to be explored. We characterized the genetic and phenotypic variation of a collection of six wild *S. sclarea* populations from Croatia, sampled along an altitudinal gradient, and of populations of three *S. sclarea* cultivars. We showed low level of genetic diversity for the two *S. sclarea* traditional cultivars used for essential oil production and for ornamental purposes, respectively. In contrast, a recent cultivar resulting from new breeding methods, which involve hybridizations among several genotypes rather than traditional recurrent selection and self-crosses over time, showed high genetic diversity. We also observed a marked phenotypic differentiation for the ornamental clary sage compared with other cultivated and wild clary sages. Instead, the two cultivars used for essential oil production, a traditional and a recent, respectively, were not phenotypically differentiated from the wild Croatian populations. Our results also featured some wild populations with high sclareol content and early-flowering phenotypes as good candidates for future breeding programs. This study opens up perspectives for basic research aiming at understanding the impact of breeding methods on clary sage evolution, and highlights interesting avenues for clary breeding programs.

## Introduction

A road-map of the genetic and phenotypic diversities in both crops and their wild-related species can help identifying valuable genetic resources for further crop breeding. Over the past century, agricultural practices caused a dramatic erosion of biodiversity in major crop plants [1]. The constant need for genetic improvement to enhance crop performance prompted the search for new genetic resources. Crop-related wild species, the wild plants closely related to cultivated crops, have a high potential genetic diversity for improvement of future breeding programs [2,3].

Despite the increasing use of medicinal aromatherapy, genetic and phenotypic variation among wild and cultivated Perfume, Aromatic and Medicinal Plants (PAMP) are still little explored (but see [4,5]). Plant specialized metabolites, such as alkaloids or terpenes, are the characteristics for which PAMP are valued. Traditional phytochemical studies have invested massive effort to isolate and characterize these chemical components, but only on a handful of elite samples. In nature, PAMP chemical composition showed a huge variation which represents a potential untapped allelic variation for breeding programs. However, the plant specialized metabolite composition of a plant depends on both environmental and genetic factors [8,9]. For instance, altitude potentially influences plant specialized metabolism as a combination of abiotic and biotic factors: temperature, hygrometry, wind speed and water availability change with altitude, promoting the development of distinct microbial and insect communities [10,11]. For targeted genetic enhancement of PAMP performances, breeders need to tease apart the environment and genetic effects at the origin of PAMP chemical variation. Common garden experiments including several genotypes sampled along an altitudinal gradient can help identifying interesting phenotypic variation that has only a genetic origin. Interesting phenotypes can then be selected for PAMP breeding programs in order to increase specialized metabolites yield. Common garden experiments can also be coupled with the characterization of the genetic variation at candidate genes involved in the specialized metabolite production pathways to point out interesting genetic variation, or lack of variation due to recent selection imposed by human. The chemical variation in crop and wild PAMP, and the underlying genes involved, are therefore good targets for breeding programs, but have generally not received a great deal of attention. Genetic study coupled with a phenotypic characterization of chemical variation of the crop and wild PAMP populations are therefore needed.

The clary sage, *Salvia sclarea* L., is a biennial or short-lived perennial with a native distribution in Northern Mediterranean, but has a vast geographic distribution towards Central Asia and the Middle East [12,13], and also become a weed outside its native range including North America. Clary sage has been traditionally used as a medicinal plant for its antispasmodic, carminative and estrogenic properties [14], and it is now mainly used for its flagrance. Clary sage essential oil, appreciated for its herbaceous and earthy scent, indeed enters in the composition of numerous high-end fragrances. Currently, several clary sage cultivars exist for different uses, *i.e.*, oil production or ornamental purposes, and result from traditional or recent breeding methods. Traditional clary sage breeding methods consist of successive cycles of selection and inbreed crosses among individuals from the same population sharing close candidate phenotypes. For oil production, candidate phenotypes show high sclareol and linalool content and, when possible, early flowering. Indeed, the most valued compounds of clary sage is Sclareol, a diterpene extracted from clary sage straw, used as starting material for the synthesis of Ambroxide^®^, characterized by a musky odor and scent fixative ability, and linalool, a monoterpene and its acetylated derivative linalyl acetate [12–16]. Early flowering is also a good candidate breeding trait in clary sage. It is widely assumed that the productivity of late-flowering clary sage plants is negatively impacted by summer drought. This rational is borrowed from breeding programs of chamomile, a medicinal plant which also produces valuable terpenoids in flowers [5]. Thereby, the traditional clary sage breeding led to the creation of cultivars used for essential oil production, *e.g.*, the “Milly” cultivar. Clary sage used for oil production has pink to purple flowers, a phenotype which stands in stark contrast with cultivars used for ornamental purposes which shows white flowers, *e.g.*, the “Vatican White” cultivar. More recently, the raising global demand for sclareol in countries where clary sage is cultivated, including Hungary, Bulgaria, Moldavia, Romania and France, has encouraged new breeding initiatives. The recent clary sage breeding approach involves recurrent selection with successive cycles of three steps: intermating between best-performing populations, evaluation of traits of interest, and selection of best-performing individuals. This new breeding method that involves the mixing of several best performing populations led to the creation of a clary sage variety with high sclareol yield, named “Scalia”. However, the “Scalia” variety displays a late-flowering phenotype, an unwanted phenotype. Further recurrent selection cycles led to the development of the “Toscalia” variety, which is characterized by a high sclareol yield without a late-flowering phenotype. While the Toscalia has an attractive phenotype for breeders, it is still not flowering early enough. Breeders are therefore still seeking for candidate populations with early-flowering, and if possible higher sclareol and linalool contents. Wild clary sage populations represent a potential source of untapped allelic variation. However, phenotypic variation for flowering time, sclareol and linalyl acetate contents of *S. sclarea* cultivars used for different purposes, as well as of wild clary sage populations, are still lacking. The genetic relationships among cultivated and wild clary sages, and their respective genetic diversity, also remained unknown.

We investigated genetic and phenotypic variation among populations of three clary sage cultivars (two traditional cultivars, the Milly and the White Vatican, and one recent cultivar, the Toscalia) and wild Croatian clary sage populations. The wild clary sage populations were sampled along an altitudinal gradient to screen for potential untapped phenotypic and genetic variation for future breeding programs. We chose Croatia because wild clary sage is widespread in this country and there is no cultivation of clary sages there, avoiding the sampling of feral clary sages that may bias the interpretations of our results. Using a genetic dataset (Internal transcribed Spacer 1 marker, referred as to “ITS” hereafter, and two candidate genes involved in the diterpene production pathways [17]), we answered the following questions: do genes involved in the sclareol production pathways show high diversity among wild and cultivated clary sage populations or signature of recent selection? More generally, what is the genetic diversity of wild and cultivated clary sages? What are the genetic relationships among cultivated and wild clary sages? Second, we grew in a common environment in France the wild and cultivated clary sages and characterized their phenotypic variation for sclareol and linalyl acetate contents, as well as their flowering time. We answered the following questions: Is there natural variation in flowering time and sclareol contents among wild and cultivated populations, and among wild populations? Our study had a dual objective: orientating breeding towards promising strategies to boost sclareol yield, and improving our general understanding of the impact of breeding methods of the evolution of a perfume, aromatic and medicinal plant.

## Material and Methods

### Plant material

During summer 2016, seeds from 60 wild *S. sclarea* individuals were collected at six sites in Croatia (Table 1 and Figure 1). These six sites were chosen to represent an altitudinal gradient: Makarska (Coast, altitude=80 m, *N*=10 individuals), Blato (Coast, 80 m, *N*=10), Zaostrog (Coast, 40 m, *N*=10), Kljenak (mountains, 360 m, *N*=10), Ravca (mountains, altitude=550 m, *N*=10), Banja (plateau, 80m, *N*=10). We chose Croatia because wild clary sage is widespread in this country and there is no cultivation of clary sages there. We therefore avoided the potential sampling of feral populations. Seeds were stored at 4 °C and 40% humidity until sowing. We also obtained seeds of cultivated *S. sclarea* from the “Vatican White”, “Toscalia” and “Milly” cultivars. Note that under the “cultivar” umbrella fall cultivated individuals resulting for several crosses between individuals sharing close phenotypes for sclareol and linalool contents, flowering time and flower color. As explained in the introduction, the Milly cultivar is traditionally used for perfume and aromatherapy purposes, whereas the Vatican White population is propagated and commercialized for ornamental purposes. The Toscalia variety is originated from recent breeding for commercial purpose for essential oil. Toscalia and Milly seeds were kindly donated by the French National Conservatory of Perfume, Medicinal, Aromatic and Industrial Plants (CNPMAI) and the French Interprofessional Institute of Perfume, Medicinal, Aromatic and Industrial Plants (ITEIPMAI), respectively. There, the Milly cultivar is maintained as a population of clary sage that are recurrently inter-bred. *Salvia sclarea ‘*Vatican white’ seeds were purchased from Jelitto (Germany). Croatian wild seeds were collected in 2017 but are not considered a strictly protected species according to the Croatian Ordinance and therefore do not fall under the Nagoya protocol. Note that the Vatican white has white flowers whereas ‘Milly’, ‘Toscalia’ and wild *S. sclarea* studied have different shades of purple-pink.

**Table 1.**
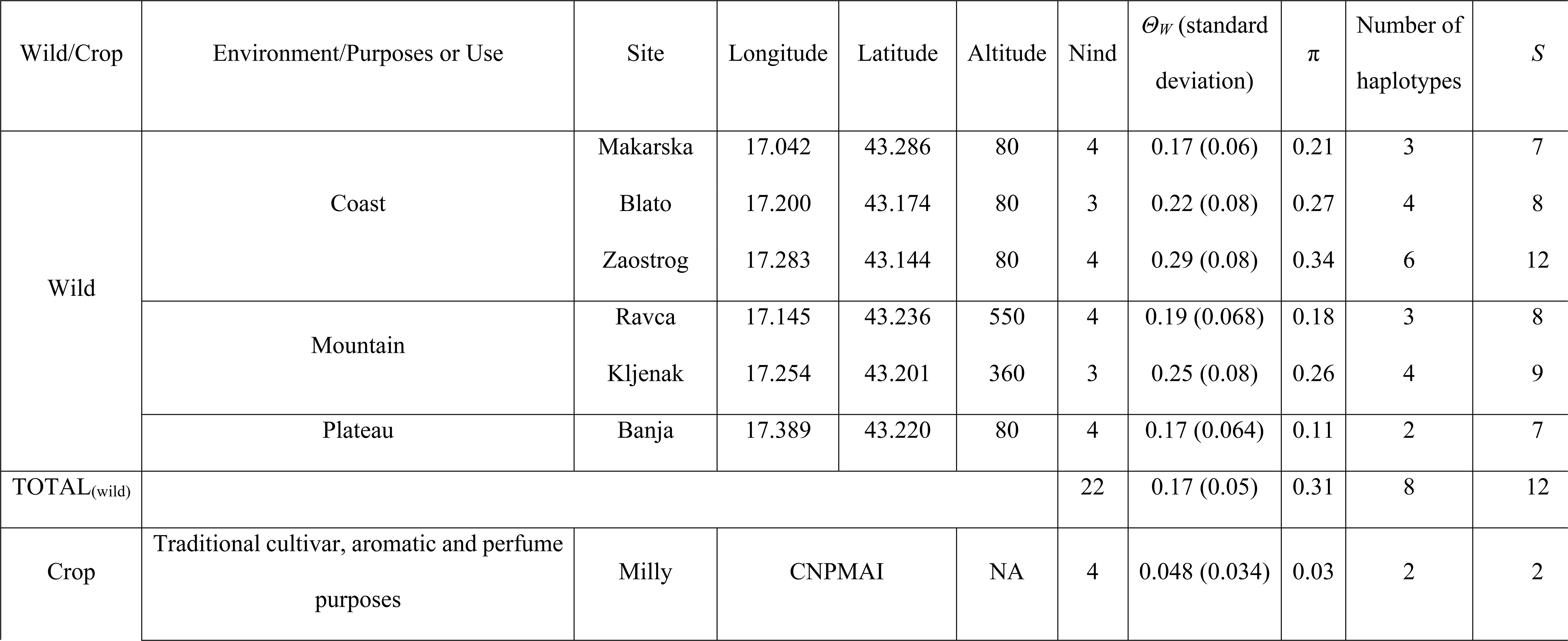
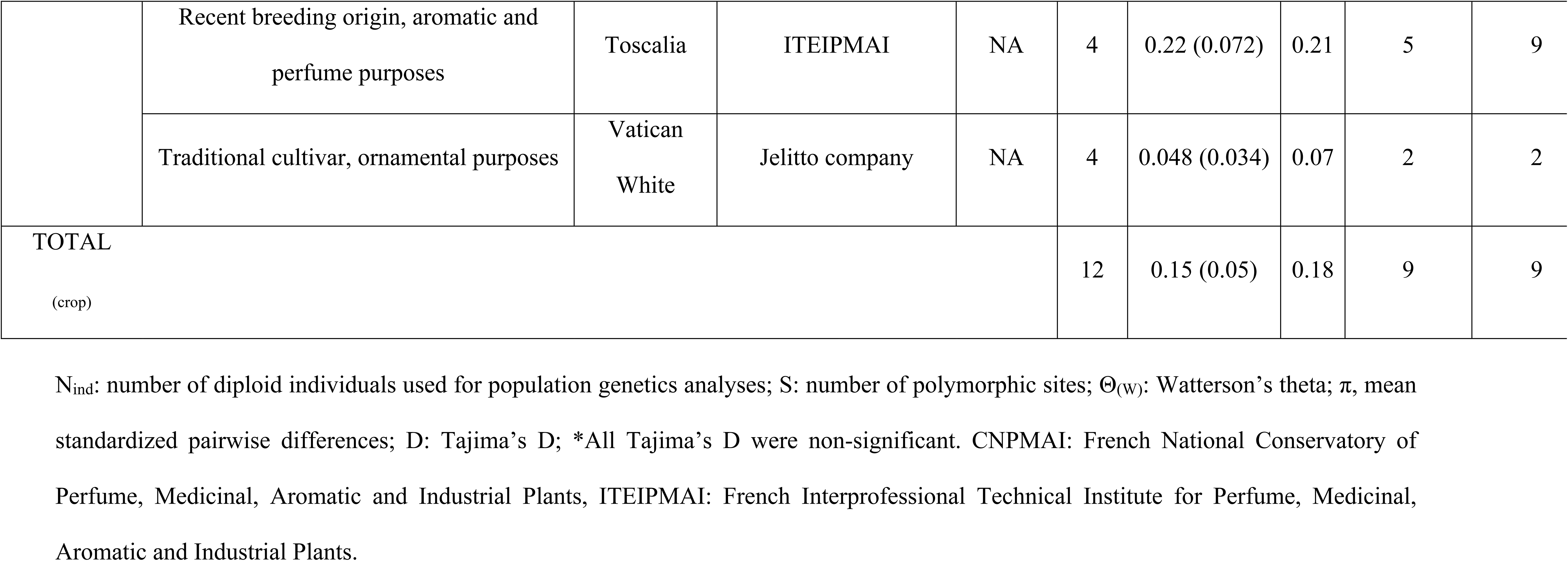
Genetic polymorphism for the wild and cultivated clary sages, *Salvia slarea*, estimated with the concatenated haploid sequences from the ITS 1, DXS2 and CMK loci (*N*=68 haploid individuals).

**Figure 1.**
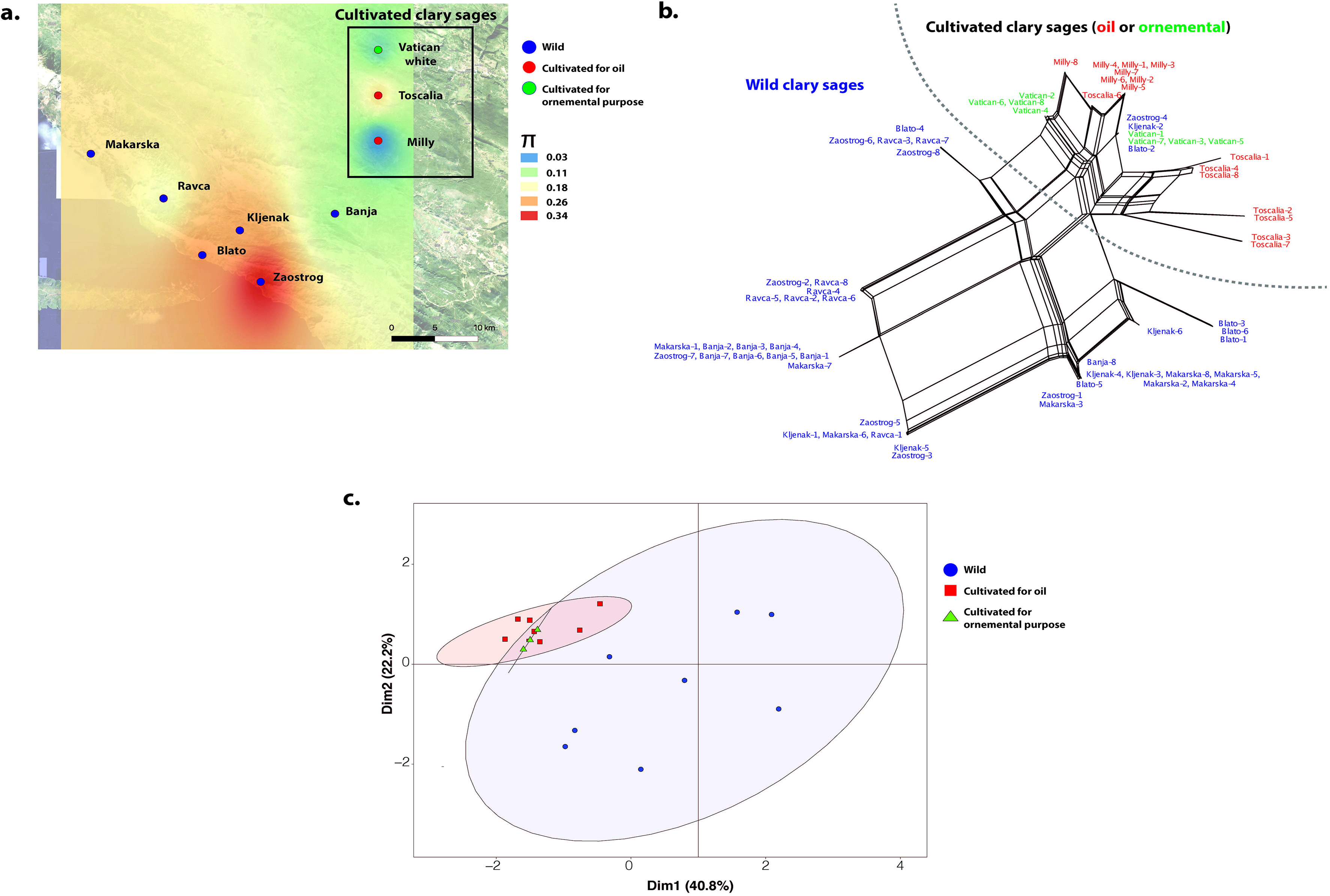
Genetic diversity and differentiation of the wild and cultivated clary sages (*Salvia sclarea*) (*N*=68 haploid individuals, nine sites, three concatenated markers CMK, ITS, DXS2). **a**. Spatial genetic diversity (π) per site for wild (six sites) and cultivated (three sites) clary sages interpolated across Croatia. The wild Croatian sites are represented by blue circles. As a matter of comparison, the three cultivar populations (red and green dots depending on their uses, for oil or ornamental purposes, respectively) were added on the map to visualize their genetic diversity relatively to the wild populations, but note they were retrieved from repository in France and Germany, and therefore their X and Y coordinates were defined arbitrarily. **b.** Splitstree of wild and cultivated clary sages. Reticulations indicate the occurrence of recombination. The color corresponds to wild (blue) and cultivated (red and green) clary sages **c.** Principal Component Analysis of the genetic variation observed between wild and cultivated sages, using the phased haploid sequences of the CMK, ITS and DSX2 markers. Dots are colored according to the wild (blue) or cultivated (red and green) individual status. Some dots overlapped as some individuals did not show any variation among each other.

### Genetic diversity and differentiation

For each sampled site for the wild *S. sclarea*, and each *S. sclarea* cultivar, three to four individuals were grown in growth chamber (16h of day; 27 °C during the day, 21 °C at night; 60% hygrometry) for six weeks. DNA was extracted from leaves using the Qiagen DNeasy 96 Plant Kit following manufacturer recommendations. Genomic fragments from *DXS2*, *CMK* and *ITS 1* loci were amplified by PCR using primers listed in Table S1. Sequences of PCR products were obtained through Sanger sequencing performed by Eurofins Genomics. Sequences were aligned using the ClustalW algorithm implemented in the MEGA7 software [18]. Heterozygous sites were visualized in Sanger sequencing chromatograms provided by Eurofins Genomics and manually annotated in MEGA7 using the IUPAC code.

We checked the neutrality of the markers to be used for population genetic analyses with *Tajima’s D* test [19]. When *Tajima’s D* test was non-significant, we assumed the sequences were evolving neutrally. However, *Tajima’s D* can be significant in case of bottleneck or expansion, regardless selection. We therefore further test for neutrality of the sequences with the McDonald and Kreitman test (MKT) [20]. The MKT was only possible when a sequence of a *Salvia* outgroup species was available on NCBI for a given marker. The resulting neutrally-evolving sequences were then concatenated. Because of the occurrence for each sequenced marker of heterozygous sites encoded with the IUPAC code, haploid sequences were phased for each individual with the PHASE algorithm implemented the DnaSP software [21]. Haploid sequences obtained (two per individual) were then used for further analyses below.

We computed *Watterson’s θ* [22] and *Tajima’s D* [19] with the DnaSP software [21] for each of the six sites for the wild *S. salvia* and each cultivar (*i.e.*, Vatican White, Toscalia and Milly). We investigated genetic relationships among wild and cultivated samples with a splitstree and a principal component analysis (PCA). For the splitstree, we used the NeighborNet distance transformation and equal angle splits transformation implemented in SplitsTree v 4 [23]. For the PCA, we used the dudi.pca function from the ade4 [24] and adegenet [25] R packages (R Development CoreTeam, URL http://www.R-project.org).

### Phenotypic characterization: field trial, flowering time measurement, and sclareol and linalyl acetate quantification by Gas chromatography–mass spectrometry (GC-MS)

Seeds from three wild clary sage sites, representing the three macro-environments (*i.e.*, Makarska-coast, Ravca-mountain and Banja-plateau, Figure 1a), and from populations of the Milly and Vatican White cultivars were sown on August 3^rd^, 2017, and then grown in a growth chamber (16h of day; 27 °C during the day, 21 °C at night; 60% hygrometry) for four months. On November 7^th^, 2017, 93 seedlings (between 15 and 23 per site or cultivar) were transplanted in the field in Chemillé (France, W 43'43" / N 47°12'36") to allow the plants overwintering. Plants were placed every 0.4 m × 0.7 m according to a randomized layout. Flowering time was defined as the date of opening of the first flower and was recorded between May 31^st^ and June 29^th^, 2018, until all plants have flowered. For calyx sampling, inflorescences were cut on July 19^th^, 2018 at the beginning of the afternoon. Mature calyces (harboring green achenes) were cut from all over the inflorescence, immediately placed in dry ice and stored at −80 °C at the end of the day until further analysis.

Clary sage calyces were immersed in hexane without grinding to extract metabolites present at the surface. For each individual, 12 mL of solvent were used to extract metabolites from eight mature calyces. The 10-undecen-1-ol was employed as internal standard and was added in the solvent before extraction at a concentration of 0.75 mM. After 2 h of agitation, 1 mL of extract was centrifuged at 13,000 rpm for 10 min to eliminate potential dust or debris. Samples were directly analyzed by GC-MS with a 7890B/5977A GC-MS system from Agilent and according to a protocol adapted from Laville et al. [26]. The system was equipped with a 10 m guard column and a Rxi-5Sil MS column (length, 30 m; inner diameter, 0.25 mm; film thickness, 0.25 μm) (Restek, Bellefonte, PA, USA). A total of 1 μl of sample was injected with a split ratio of 30:1. Oven temperature was set to 110 °C for 2 min, then increased to 270 °C at a rate of 10 °C/min, and finally set to 270 °C for 2 min. Other temperatures were set as follows: injector, 270 °C; transfer line, 270 °C; source, 230 °C; quadrupole, 150 °C. The carrier gas was helium at a constant flow of 1 mL/min. The quadrupole mass spectrometer was switched on after a solvent delay of 5 min and was programmed to scan from 35 u to 350 u. Data analysis was carried out using the MassHunter Quantitative Analysis software from Agilent. Sclareol and linalyl acetate were identified by comparison with analytical standards purchased from Sigma-Aldrich. Absolute quantification of sclareol and linalyl acetate was performed using a calibration curve prepared with analytical standards.

### Statistical analyses

In the presence of a domestication syndrome, we expect differences in phenotypes of cultivated accessions compared to wild accessions. We can also expect phenotypic differences between the cultivated clary sages depending on their uses (ornamental or oil production), and between wild *S. sclarea* plants sampled at sites along an altitudinal gradient in Croatia. Indeed, the selective pressures associated with the three environmental conditions from which the three populations were sampled (Makarska, Coast; Ravca, mountains; Banja, plateau) may have led to natural variation in phenotypes. In order to study the influence of the use of the cultivated sages and the wild status of each plant, we fitted a generalized linear model as follows:

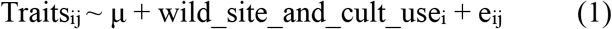

Where μ is the overall mean of the phenotypic trait (flowering time or sclareol or linalyl acetate content), ‘wild_and_cult_use’ is the fixed effect of the status of each plant (wild from Makarska/Coast, wild from Ravca/mountains, wild from Banja/plateau, cultivated clary sage used for ornamental purpose, *i.e.*, White Vatican cultivar, and cultivated clary sage used for oil production, i.e., Milly cultivar) and e is the residual. We ran a general linear model, with a residual term assumed to be normally distributed. Statistical significance of the difference between status was tested with a pairwise a Kruskal-Wallis rank test.

## Results

### Genetic diversity and relationships among cultivated and wild clary sages

Tajima’s *D* and MKT showed that the *ITS*, and the two genes involved in the diterpene production pathway, *DXS2* and *CMK*, evolved neutrally (Table S2). Table 1 gives genetic diversity estimates for each sampling site for the wild clary sages, and for the three cultivars. The map of interpolated pairwise nucleotide diversity, π, by site (Figure 1a) showed that genetic diversity was highest for the two wild coastal sites (Blato and Zaostrop) and one mountain site (Kljenak), and the lowest in the plateau site (Banja) (Table 1 and Figure 1a). The cultivated clary sages showed lower level of genetic diversity than the wild populations, except the recent Toscalia cultivar (Table 1 and Figure 1a).

The splitstree and the PCA (Figure 1, c and d) revealed a genetic differentiation between wild and cultivated clary sages. Within the cultivated clary sages, the Vatican white showed less variation than the Milly and Toscalia cultivar, and the Toscalia were more differentiated than the Milly and the White Vatican cultivars.

### Phenotypic variation between wild and cultivated clary sages, and within each group

In total, 20 plants out of the 93 initial transplanted seedlings during the winter of 2017 did not survive the winter. The 73 remaining plants (between 10 and 22 per wild site or cultivar) flowered between May 31^st^ and June 29^th^, 2018, that is, between 301 and 330 days after sowing. Flowering dates were consistent with data available in the literature; indeed, clary sage is usually reported to flower in June or July [12].

Overall, we observed significant differences in flowering time, sclareol and linalyl acetate among wild clary sages, clary sages cultivated for ornamental purposes and for oil production (Figure 2a and Table 2). For wild clary sages, no significant differences in sclareol and linalyl acetate of calyces were observed among plants coming from coast, mountain or plateau (Figure 2, c and d). Significant differences were observed for flowering time, with the Makarska/coastal plants flowering in average 5.044 days later than other wild plants (*P*=0.046, Figure 2b). When comparing the cultivated and wild plants, the Vatican White plants, used for ornamental purposes, produced significantly less linalyl acetate (30% less) than the Milly cultivar and the wild clary sage plants (Figure 2c). The Vatican White plants also produced significantly less sclareol (−0.12 mg/calyx in average, *P*<0.0001, Figure 2d) compared to the Milly cultivar and the Makarska/coast and Banja/plateau wild plants. Vatican White plants flowered significantly earlier than the Milly cultivar and the wild Makarska plants (10 days earlier in average) (Figure 2b)

**Figure 2.**
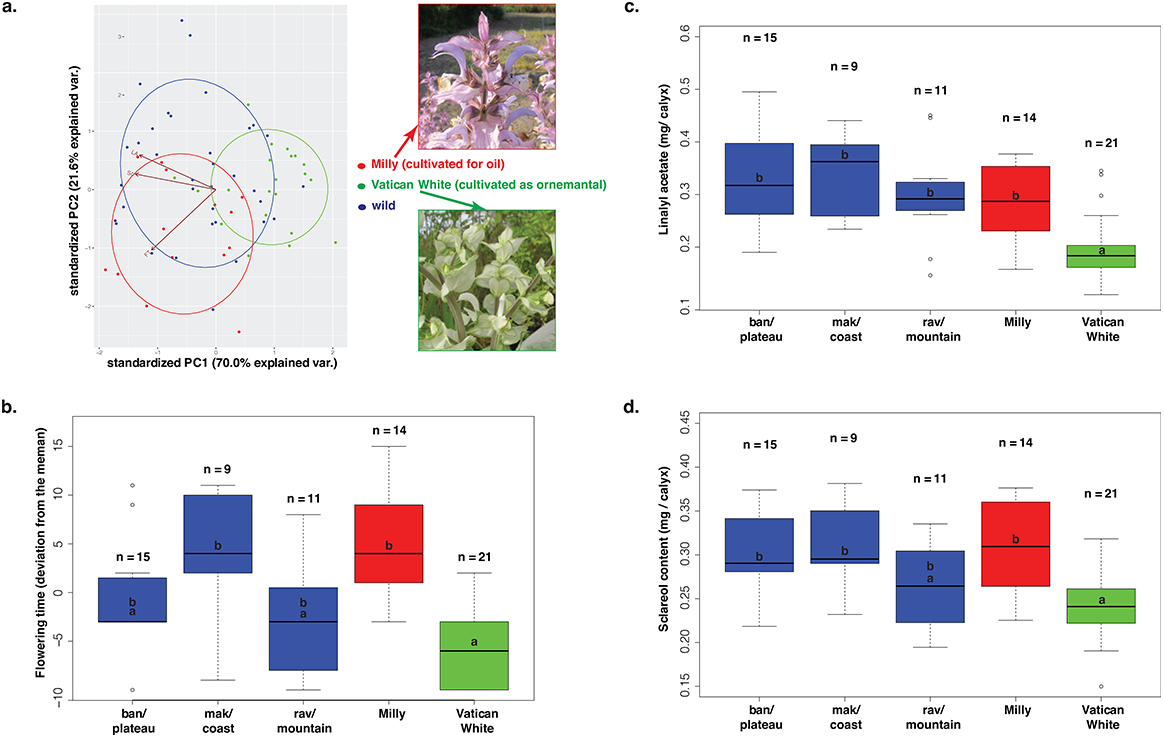
Phenotypic variation (for flowering time, sclareol and linalyl acetate contents) for the wild and cultivated clary sages (*Salvia sclarea*) measured in the common garden at Chemillé in France in 2018. a. Principal component analysis on phenotypic traits for the wild clary sages (blue), the Milly cultivar (red), and the White Vatican cultivar (green) with the photos of the flowers of representative Vatican White and Milly clary sage plants. b, c, d. Date of flowering (expressed in deviation from the mean), linalyl acetate and sclareol contents (expressed in mg/calyx), respectively, for each wild clary sage population (blue), the Milly cultivar (red), and the White Vatican cultivar (green). Numbers indicate the number of samples for each population, letters indicate the result of a Kruskal-Wallis rank test: values harboring the same letter are not significantly different

**Table 2.**
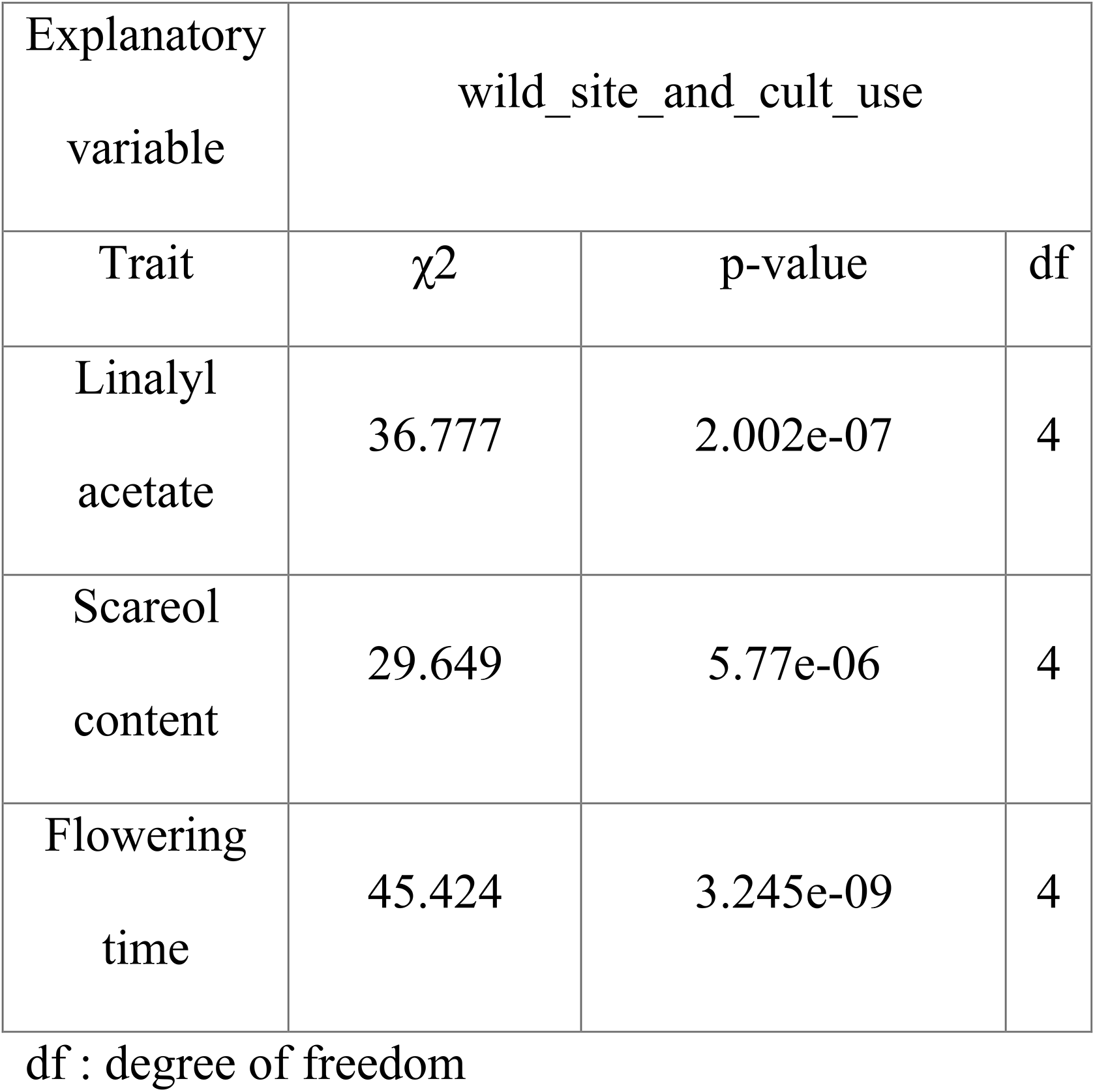
Summary of the effect of the wild site of origin and the use for cultivation of the cultivated clary sage on phenotypic trait variation (Model 1) in *Salvia sclarea*.

## Discussion

We provided a first insight into crop and wild genetic and phenotypic variation of a perfume, medicinal and aromatic plant, the clary sage. Our results showed a lower level of genetic diversity for the traditional clary sage cultivars, whatever their uses, compared with the Croatian wild populations. In contrast, the recent Toscalia cultivar, resulting from new breeding methods, showed a high genetic variation, not significantly different from the Croatian wild populations. Phenotypically, a strong divergence is observed for the ornamental clary sage compared with the two cultivars used for essential oil production. Interestingly, the cultivars used for essential oil production were not significantly differentiated from the Croatian wild populations, whatever the area/altitude of origin of the wild population. We also did not observe phenotypic variation among wild Croatian clary sage populations. Our results finally pinpoint certain Croatian wild populations with interesting sclareol and linalyl acetate contents for future clary sage breeding.

### A lower genetic diversity for the traditional clary sage cultivars, whatever their uses

We did find a lower genetic diversity for the traditional cultivars used for essential oil production (Milly) and for ornamental purposes (Vatican White). Traditional clary sage breeding methods consist of successive cycles of selection and inbreed crosses among individuals from the same population sharing close candidate phenotypes, either for sclareol content and/or early-flowering for oil production, or flower color for ornamental purposes. The recurrent selection and bottlenecks implied have likely played a role in the loss of genetic diversity observed for the two cultivars. In contrast, the recent breeding strategy consists of hybridizations of different genotypes to diversify the genetic basis for *S. sclarea* quality though raising the capacity of essential oil accumulation from the same population. Such new breeding method has resulted in the recent Toscalia cultivar, which, not surprisingly showed high level of genetic diversity, equivalent to the one found in the wild Croatian populations studied here.

The lower diversity of traditional clary sage cultivars may reflect a bottleneck during domestication but also a post-domestication erosion [27]. Further sampling of several cultivars from various geographical origins and representing a gradient of domestication, from old and traditional, to recent cultivars, is now required to widen our conclusions. Besides, small populations that underwent bottlenecks thought recurrent selection during domestication can also be expected to show evidence for recent selection in genes involved in key agronomic traits [28]. Here, we did not find any signatures of selection in two genes involved in chemical production pathways, the CMK and DMSX2 genes. Selection could have played a role in other genes involved in the production pathways that were not sequenced in our study, or/and our sampling was too limited. High-throughput sequencing of several varieties with various phenotypes, and association between phenotypes and genotypes [29,30], or scan for selective sweeps along the cultivated and wild clary sage genomes [31], could provide deeper insights in the genomic processes involved during the domestication of clary sage.

### Phenotypic divergence of the cultivated clary sages depending on their uses, and lack of phenotypic differentiation among wild Croatian clary sages

We showed a strong phenotypic divergence for the ornamental clary sage cultivar, with an earlier flowering and much lower sclareol and linalyl acetate content than the wild Croatian clary sage populations and the cultivar used for essential oil production. Indeed, Vatican White plants flowered 10 days earlier and produced 20% less sclareol and 30% less linalyl acetate than the cultivar used for oil production. In addition, Vatican White cultivar has light flowers that stand in stark contrast with the flowers of other cultivars used for oil production (*e.g.*, Milly or Toscalia), or of *S. sclarea* wild populations, that display darker shades of pink and purple. In other plant species, the pink or purple color of flowers is often due to the presence of specialized metabolites belonging to the flavonoid family [32]. Traditional breeding methods have therefore implied simultaneous decrease in flavonoid biosynthesis and terpenoid biosynthesis at the origin of the low oil content and different flower color. If this downregulation is operated at transcriptome level, it would be relevant to conduct a comparative transcriptomic study of Milly and Vatican White flower organs. A differential expression of biosynthetic genes would validate the initial hypothesis, and differentially expressed transcription factors would be good candidates for an involvement in specialized metabolism regulation or flowering time control. Since sclareol and linalyl acetate are mainly produced by glandular trichomes, the lower metabolite content observed in Vatican White plants may also be due to a lower glandular trichome density. In *Artemisia annua*, the amount of artemisinin found in leaves is tightly correlated to glandular trichome density [33]. Careful measurements of glandular trichome density are needed to test this hypothesis.

In contrast, we showed a lack of variation in metabolite content between the cultivated clary sage used for oil production and the wild clary sage populations. This lack of crop-wild variation for metabolite content suggests that it may not be a trait to pick up in the Croatian wild populations for future breeding programs. Instead, we did observed variation for flowering time among wild clary sage populations. In particular, the Ravca population showed early-flowering phenotypes. This population could be a good target for future breeding program of clary sage used for essential oil production because of their early flowering and relatively high sclareol content, close to the metabolite content found in the cultivar used for essential oil production (Milly).

Finally, planted in a common environment, the six wild populations sampled across an environmental gradient did not show significant differences in metabolite content. This result suggests no genetic difference for metabolic contents among wild populations. Therefore, while plants from the wild Croatian populations can be used for breeding programs as a whole regarding their metabolic content, the Ravca population can be used specifically for its early flowering. Additional common gardens including several wild and cultivated clary sage populations in different sites across an altitudinal gradient are now required to guide future clary sage breeding programs. Specific environmental conditions may induce variation yield and chemical composition in metabolite contents, as previously shown in a related species, *Salvia officinalis* [34].

## Conclusions

Clary sage produces sclareol, a terpenoid widely used in perfume industry for the hemisynthesis of ambroxide, a valued perfume component with remarkable fixative properties. In response to the development of ambroxide use in functional perfumery, the global demand for sclareol currently raises, prompting efforts aiming at increasing sclareol supply. Breeding programs aims at increasing the yield of sclareol production from clary sage through targeted genetic improvement. The purpose of our study was to enhance knowledge on phenotypic and genetic variation of wild and cultivated clary sage to provide a first basis for clary sages breeding programs. A larger number of markers and samples of wild and cultivated sages (used for ornamental and oil purposes) across the clary sage distribution is however now required to better understand the clary sage evolution, and provide further directions for breeding programs.

## Supplementary Materials

Table S1. Primers used to amplify genomic fragments from DXS2, CMK and ITS loci to characterize the genetic diversity of *Salvia sclarea*

Table S2. Summary statistics obtained for the three markers (ITS, CMK, DXS2) for wild and cultivated *Sclarea salvia*,

## Author Contributions

Conceptualization, ABo, ABe and MD; methodology, ABo, AC, SD, MD, FG, CCho and CCha; software, AC; validation, CCha and CCho; formal analysis, CCha, CCho, AC, SD, FG; investigation, AC, CCha, SD, CCho, and FG; resources, MD and EMS; data curation, CCha, SD, FG, AC, ABo; writing—original draft preparation, AC, CCha, SD, MD; writing—review and editing, AC, SD, ABo, ABe, MD; supervision, ABo, ABe, MD and SD; funding acquisition, ABo, ABe and MD. All authors have read and agreed to the published version of the manuscript.

## Acknowledgments

We thank Bernard Pasquier, former Director of CNPMAI, France, and Jean-Pierre Bouverat-Bernier, former Director of Iteipmai, France, for kindly giving us 'Milly-la-Forêt' and 'Toscalia' accessions respectively. We also thank Clémentine Fayol, Alan Walton and Guillaume Frémondière for growing and phenotyping plants in ITEIPMAI, France. We are grateful for good advice and connexions of Sonya Jakovlev, Paris-Saclay University, France. To finish, thanks to Dubravka Stepić, from the Croatian Ministry of Environment and Energy, to authorize the sampling of wild clary sage in Croatia and for the fruitful discussions about the Nagoya protocol. This research was funded by Plant Biology and Breeding department of INRAE, the French National Research Agency (ANR), and the grants Program LabEx Saclay Plant Sciences-SPS (ANR-10-LABX-40-SPS).

## References

1. Kilian B, Graner A. NGS technologies for analyzing germplasm diversity in genebanks. Briefings in Functional Genomics. 2012;11: 38–50. doi:10.1093/bfgp/elr046

2. Zhang H, Mittal N, Leamy LJ, Barazani O, Song B-H. Back into the wild—Apply untapped genetic diversity of wild relatives for crop improvement. Evolutionary Applications. 2017;10: 5–24. doi:10.1111/eva.12434

3. Castañeda-Álvarez NP, Khoury CK, Achicanoy HA, Bernau V, Dempewolf H, Eastwood RJ, et al. Global conservation priorities for crop wild relatives. Nature Plants. 2016;2: 16022. doi:10.1038/nplants.2016.22

4. Raymond O, Gouzy J, Just J, Badouin H, Verdenaud M, Lemainque A, et al. The Rosa genome provides new insights into the domestication of modern roses. Nature Genetics. 2018;50: 772–777. doi:10.1038/s41588-018-0110-3

5. Otto LG, Mondal P, Brassac J, Preiss S, Degenhardt J, He S, et al. Use of genotyping-by-sequencing to determine the genetic structure in the medicinal plant chamomile, and to identify flowering time and alpha-bisabolol associated SNP-loci by genome-wide association mapping. BMC Genomics. 2017;18: 1–18. doi:10.1186/s12864-017-3991-0

6. Meyer RS, DuVal AE, Jensen HR. Patterns and processes in crop domestication: an historical review and quantitative analysis of 203 global food crops. New Phytologist. 2012;196: 29–48. doi:10.1111/j.1469-8137.2012.04253.x

7. Glémin S, Bataillon T. A comparative view of the evolution of grasses under domestication. New Phytol. 2009;183: 273–290.

8. Figueiredo AC, Barroso JG, Pedro LG, Scheffer JJC. Factors affecting secondary metabolite production in plants: volatile components and essential oils. Flavour and Fragrance Journal. 2008;23: 213–226. doi:10.1002/ffj.1875

9. Moore BD, Andrew RL, Külheim C, Foley WJ. Explaining intraspecific diversity in plant secondary metabolites in an ecological context. New Phytologist. 2014;201: 733–750. doi:10.1111/nph.12526

10. Mahzooni-Kachapi SS, Mahdavi M, Jouri MH, Akbarzadeh M, Roozbeh-nasira’ei L. The effects of altitude on chemical compositions and function of essential oils in stachys lavandulifolia vahl. (Iran). International Journal of Medicinal and Aromatic Plants. 2014;4: 107–116.

11. Foutami IJ, Mariager T, Rinnan R, Barnes CJ, Rønsted N. Hundred Fifty Years of Herbarium Collections Provide a Reliable Resource of Volatile Terpenoid Profiles Showing Strong Species Effect in Four Medicinal Species of Salvia Across the Mediterranean. Frontiers in Plant Science. 2018;9: 1877. doi:10.3389/fpls.2018.01877

12. Wagner S, Pfleger A, Mandl M, Böchzelt H. Changes in the Qualitative and Quantitative Composition of Essential Oils of Clary Sage and Roman Chamomile During Steam Distillation in Pilot Plant Scale. Distillation - Advances from Modeling to Applications. 2012; 141–158.

13. Nasermoadeli S, Rowshan V. Comparison of Salvia sclarea L. Essential Oil Components in Wild and Field Population. International Journal of Agriculture and Crop Sciences. 2013;4: 1997–2000.

14. Lattoo SK, Dhar RS, Dhar AK, Sharma PR, Agarwal SG. Dynamics of essential oil biosynthesis in relation to inflorescence and glandular ontogeny in Salvia sclarea. Flavour and Fragrance Journal. 2006;21: 817–821. doi:10.1002/ffj.1733

15. Schmiderer C, Grassi P, Novak J, Weber M, Franz C. Diversity of essential oil glands of clary sage (Salvia sclarea L., Lamiaceae). Plant Biology. 2008;10: 433–440. doi:10.1111/j.1438-8677.2008.00053.x

16. Sharopov FS, Setzer WN. The essential oil of salvia sclarea L. from Tajikistan. Records of Natural Products. 2012;6: 75–79.

17. Tholl D. Biosynthesis and Biological Functions of Terpenoids in Plants. Advances in biochemical engineering/biotechnology. 2015;123: 127–141. doi:10.1007/10

18. Kumar S, Stecher G, Li M, Knyaz C, Tamura K. MEGA X: Molecular Evolutionary Genetics Analysis across Computing Platforms. Mol Biol Evol. 2018;35: 1547–1549. doi:10.1093/molbev/msy096

19. Tajima F. Statistical method for testing the neutral mutation hypothesis by DNA polymorphism. Genetics. 1989;123: 585–595.

20. McDonald JH, Kreitman M. Adaptive protein evolution at the Adh locus in Drosophila. Nature. 1991;351: 652–654. doi:10.1038/351652a0

21. Rozas J, SÃ¡nchez-DelBarrio JC, Messeguer X, Rozas R. DnaSP, DNA polymorphism analyses by the coalescent and other methods. Bioinformatics. 2003;19: 2496–2497. doi:10.1093/bioinformatics/btg359

22. Watterson GA. On the number of segregating sites in genetical models without recombination. Theor Popul Biol. 1975;7: 256–276. doi:10.1016/0040-5809(75)90020-9

23. Huson DH, Bryant D. Application of phylogenetic networks in evolutionary studies. Mol Biol Evol. 2006;23: 254–267. doi:10.1093/molbev/msj030

24. Dray S, Dufour A-B. The ade4 Package: Implementing the Duality Diagram for Ecologists. Journal of Statistical Software; Vol 1, Issue 4 (2007). 2007. doi:10.18637/jss.v022.i04

25. Jombart T, Ahmed I. adegenet 1.3-1: new tools for the analysis of genome-wide SNP data. Bioinformatics. 2011;27: 3070–3071. doi:10.1093/bioinformatics/btr521

26. Laville R, Castel C, Filippi JJ, Delbecque C, Audran A, Garry PP, et al. Amphilectane diterpenes from Salvia sclarea: Biosynthetic considerations. Journal of Natural Products. 2012;75: 121–126. doi:10.1021/np2004177

27. Allaby RG, Ware RL, Kistler L. A re-evaluation of the domestication bottleneck from archaeogenomic evidence. Evolutionary Applications. 2019;12: 29–37. doi:10.1111/eva.12680

28. Purugganan MD, Fuller DQ. The nature of selection during plant domestication. Nature. 2009;457: 843–848.

29. Brachi B, Morris GP, Borevitz JO. Genome-wide association studies in plants: the missing heritability is in the field. Genome Biology. 2011;12: 232. doi:10.1186/gb-2011-12-10-232

30. Huang X, Han B. Natural Variations and Genome-Wide Association Studies in Crop Plants. Annual Review of Plant Biology. 2014;65: 531–551. doi:10.1146/annurev-arplant-050213-035715

31. Nielsen R. Molecular Signatures of Natural Selection. Annu Rev Genet. 2005;39: 197–218. doi:10.1146/annurev.genet.39.073003.112420

32. Tanaka Y, Sasaki N, Ohmiya A. Biosynthesis of plant pigments: anthocyanins, betalains and carotenoids. The Plant Journal. 2008;54: 733–749. doi:10.1111/j.1365-313X.2008.03447.x

33. Xiao L, Tan H, Zhang L. Artemisia annua glandular secretory trichomes: the biofactory of antimalarial agent artemisinin. Science Bulletin. 2016;61: 26–36. doi:10.1007/s11434-015-0980-z

34. Russo A, Formisano C, Rigano D, Senatore F, Delfine S, Cardile V, et al. Chemical composition and anticancer activity of essential oils of Mediterranean sage (Salvia officinalis L.) grown in different environmental conditions. Food and Chemical Toxicology. 2013;55: 42–47. doi:10.1016/j.fct.2012.12.036

35. He J, Zhao X, Laroche A, Lu Z-X, Liu H, Li Z. Genotyping-by-sequencing (GBS), an ultimate marker-assisted selection (MAS) tool to accelerate plant breeding. Frontiers in Plant Science. 2014;5: 484. doi:10.3389/fpls.2014.00484

