## Supplementary file for "Study of the genetic and phenotypic variation among wild and cultivated clary sages provides interesting avenues for breeding programs of a perfume, medicinal and aromatic plant"

**Table S1. Primers used to amplify genomic fragments from DXS2, CMK and ITS loci to characterize the genetic diversity of *Salvia sclarea*.**

| Target | Primer | Sequence |
|---|---|---|
| DXS2 | DXS2_F | 5'-GCAGTTTCTTGCCATTGCTCC-3' |
| | DXS2_R | 5'-TAATACGTACCTGGTGACCC-3' |
| CMK | CMK_F | 5'-CGAGAGGTACAGGTGGAGGA-3' |
| | CMK_R | 5'-CAAGAGTGGTCGGGTGAGAT-3' |
| ITS 1 | ITS_F | 5'-GCATCGATGAAGAACGTAGC-3' |
| | ITS_R | 5'-TCCTCCGCTTATTGATATGC-3' |

**Table S2**. Summary statistics obtained for the three markers (ITS, CMK, DXS2) for wild and cultivated *Salvia sclarea*.

| | | | | | | | | | | Coding region* | | |
|---|---|---|---|---|---|---|---|---|---|---|---|---|
| Marker | Length (base pairs) | Nseq | $N_{ind}$ | S | $\pi$ | Tajima's D | Fu and Li's D | Fu and Li's F | MacDonald Kreitman test | ORF1 | ORF 2 | ORF3 |
| ITS | 323 | 68 | 38 | 3 | 0.0415 | 2.19 ** | 0.86 NS | 1.49 NS | NS (outgroup MK124723.1 *Salvia splendens* and MF543806.1 *Salvia aethiopis*) | 1-230 | 135-323 | NA |
| CMK | 366 | 68 | 34 | 9 | 0.00699 | 0.96 NS | 1.339 NS | 1.43 NS | Not possible to find outgroups | 1-108 | 164-208 | 279-365 |
| DXS2 | 434 | 68 | 34 | 4 | 0.00398 | 2.27 ** | 0.97 NS | 1.61 NS | NS (outgroup used MK067342.1/1-280 *Salvia officinalis*) | 94-216 | 116-262 | 290-421 |

Nind: number of individuals, *S*: number of polymorphic sites, $\pi$*:* mean standardized pairwise, differences. * *Coding regions* defined with default parameters on: https://www.ncbi.nlm.nih.gov/orffinder/
